# PQBP1 couples HIV-1 capsid recognition to cGAS recruitment through conformational remodeling

**DOI:** 10.64898/2026.07.07.737097

**Authors:** Dale S Allen, Asit Manna, Christian Michel Beusch, Tianhao Zhang, Barbie K Ganser-Pornillos, Juliana Piacentini, Seolhwa Oh, Savni Prabhu, Ayush Mehta, David Gordon, Owen Pornillos, Sunnie M Yoh, Sumit K Chanda

## Abstract

Pattern recognition receptors (PRRs) must selectively engage pathogen-derived signals to potentiate inflammation and antimicrobial responses. Polyglutamine-binding protein 1 (PQBP1) acts upstream of cyclic GMP-AMP synthase (cGAS) during HIV-1 infection through the recognition of viral capsid. However, the mechanism by which capsid binding enables cGAS recruitment remains unclear. As an intrinsically disordered protein, PQBP1 samples an ensemble of conformational states whose distribution depends on ligand engagement. Here we show that capsid engagement shifts this population toward a conformation competent for cGAS binding. Capsid binding at the N-terminus of PQBP1 redistributes conformational sampling within the WW domain and distal polar-rich domain (PRD). Alanine substitutions within these capsid-responsive regions retain capsid binding yet disrupt infection-dependent cGAS recruitment, indicating that capsid binding and cGAS recruitment depend on distinct regions of PQBP1. Together, these findings define a mechanism by which HIV-1 capsid engagement remodels PQBP1 into a cGAS-competent state, coupling capsid recognition to innate immune activation.

**IMPORTANCE:** Polyglutamine-binding protein 1 (PQBP1) initiates innate immune detection of HIV-1 by recognizing the incoming viral capsid. We show that capsid engagement reshapes PQBP1 conformational dynamics, and we identify distal regions required for infection-dependent cGAS association. These findings provide a mechanistic framework for how pathogen recognition is coupled to downstream innate immune activation.

## INTRODUCTION

Innate immune responses rely on pattern recognition receptors (PRRs) to detect molecular features associated with pathogens or cellular damage while avoiding aberrant activation by self-derived ligands (1). Cyclic GMP-AMP synthase (cGAS) is a central mediator of type I interferon responses to cytosolic DNA (2). However, cGAS activation during HIV-1 infection presents a unique challenge, as viral DNA is transient, low-abundance, and shielded within an intact capsid for much of the viral life cycle (3–5). How cGAS is selectively activated during HIV-1 infection despite the limited accessibility of viral DNA is a central unresolved question in HIV innate immunity.

More broadly, many innate immune signaling pathways require cofactors that license downstream activation only under specific biological contexts. Toll-like receptor 4 depends on the co-receptor, MD-2, to detect bacterial lipopolysaccharide (6), and RIG-I requires TRIM25-dependent signaling for viral RNA recognition (7). Polyglutamine-binding protein 1 (PQBP1) is an essential upstream factor that licenses cGAS activation during acute HIV-1 infection in monocyte-derived cells (8). Previous work established that PQBP1 associates with intact HIV-1 capsids and promotes cGAS engagement. Through this mechanism, capsid recognition precedes and enables productive cGAS recruitment, allowing detection of viral nucleic acids that would otherwise escape immune surveillance (9, 10). A recent study further defines the molecular basis of this initial step, showing that the PQBP1 N-terminus engages the arginine-rich central pore of the HIV-1 capsid hexamer through charge complementation (11).

How capsid engagement is converted into productive cGAS recruitment remains unclear. PQBP1 is a multidomain intrinsically disordered protein comprising an N-terminal capsid-binding region, a WW domain, a polar-rich domain (PRD), and a C-terminal tail (Ct) (12–14). Intrinsically disordered proteins frequently regulate signaling through ligand-coupled conformational changes that expose or reshape functional surfaces distal from the primary binding interface, a process commonly described as dynamic coupling (15).

Prior biophysical studies indicate that PQBP1 undergoes ligand-responsive conformational changes outside immune contexts. Within the spliceosome complex, WBP11 binding to the PQBP1 WW domain allosterically weakens U5-15kD binding at the PQBP1 C-terminus, illustrating long-range coupling across PQBP1. Additionally, disease-associated WW domain mutations alter PQBP1 folding and its interaction with spliceosome complex (16, 17). Together, these observations establish PQBP1 as a conformationally dynamic protein whose states respond to ligand engagement, raising the possibility that HIV-1 capsid binding reshapes PQBP1 to promote cGAS recruitment. Here, we define the structural consequences of capsid binding across PQBP1 and link these changes to infection-dependent cGAS recruitment in cells.

## RESULTS

### Regions outside CA hexamer binding domain regulates PQBP1-CA hexamer interaction

Isothermal titration calorimetry (ITC) of PQBP1 truncation constructs revealed that regions outside the N-terminus substantially alter the capsid hexamer binding profile (Fig. 1A). Consistent with previous work and modeling (Fig. S1A) (11), the isolated N-terminal segment (residues 1-46) bound the capsid hexamer with high affinity in ITC experiments (Kd ∼30 nM; Figs. 1B, C). In contrast, full-length PQBP1 exhibited substantially weaker binding (K_d_ ∼6.2 µM), with apparent Kd increasing progressively across the WW-domain constructs (1-88, ∼2.5 µM; 1-104, ∼4.2 µM) (Figs. 1B, C). This length-dependent loss of affinity suggests that the WW domain and flanking regions alter how PQBP1 engages capsid. Each construct bound a single hexamer (1:1 stoichiometry) with a favorable entropic contribution (Fig. 1C). The stoichiometry indicates this affinity change is a result of one binding event rather than a change in the defined binding mode (11). The favorable entropy could reflect desolvation of the complex or conformational changes in PQBP1 or the hexamer. Together with the 1:1 stoichiometry, this suggests that a single hexamer alters PQBP1 conformational states through an entropy-dominated interaction.

**Figure 1.**
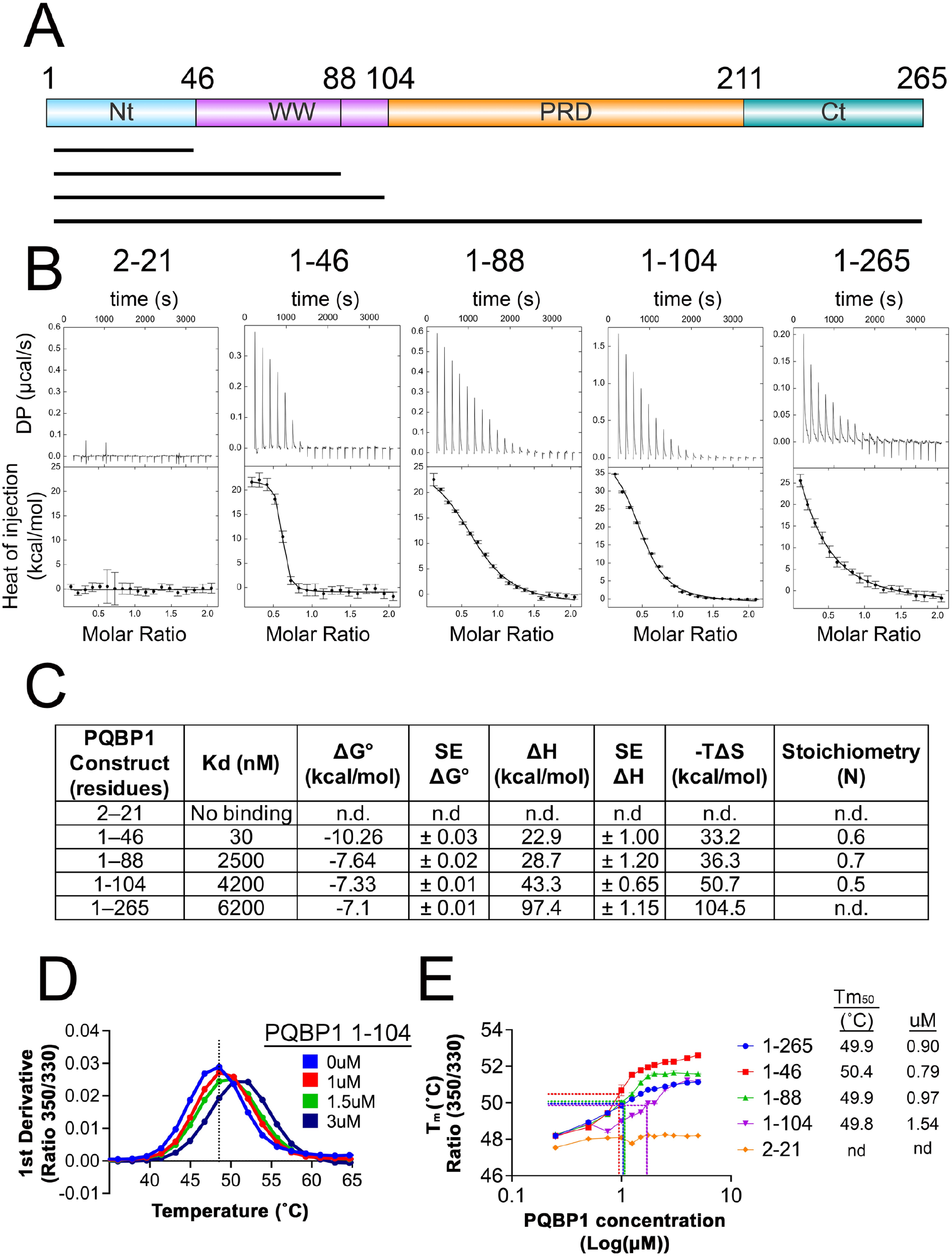
Regions outside PQBP1 Nt regulate hexamer binding (A) Domain schematic scaled to full-length PQBP1, highlighting the N-terminal (Nt), WW, polar-rich (PRD), and C-terminal (Ct) regions. Bars below indicate the truncation constructs used: 2-21, 1-46, 1-88, 1-104, and full-length (FL) 1-265. Created with IBS 2.0. (B) Representative isothermal titration calorimetry (ITC) thermograms and binding isotherms for the titration of HIV-1 capsid hexamer with PQBP1 FL and indicated truncation constructs. Upper panels: raw heat per injection; lower panels: integrated heats vs. molar ratio with fitted curves. Values are fit ± SE per construct; representative of two independent experiments. See Methods for details. (C) Thermodynamic parameters (Kd, ΔH, stoichiometry N) from ITC fits for PQBP1 FL and truncation constructs binding the HIV-1 capsid hexamer. Stoichiometry for full-length PQBP1 (1-265) was not determinable from the shallow binding isotherm at 6.2 µM; see Methods. (D) Nano-differential scanning fluorimetry (nanoDSF) thermal denaturation profiles of HIV-1 capsid hexamer (2 µM) with increasing concentrations of PQBP1 (1-104), shown as the first derivative curve of the 350/330 nm fluorescence ratio versus temperature. Representative of at least two independent experiments. (E) NanoDSF-derived apparent melting temperatures (Tm) of HIV-1 capsid hexamer upon titration of PQBP1 FL or indicated truncation constructs. Hexamer alone, Tm ∼48 °C; with full-length PQBP1, Tm ∼51 °C at saturation. Dotted lines denote the Tm₅₀ for each construct, defined as the temperature at the midpoint of the concentration-dependent Tm transition. Tm₅₀ values (°C) and apparent dissociation constants are tabulated at right. Representative of at least two independent experiments.

Nano-differential scanning fluorimetry (nanoDSF) provided additional support that the WW domain (residues 47-104) captures the majority of the capsid interaction behavior observed for full-length PQBP1. Full-length PQBP1 produced a binding profile, measured as hexamer melting temperature (Tm) versus PQBP1 concentration, intermediate between the 1-88 and 1-104 constructs (Figs. 1D, E, S1B). All constructs reached a defined Tm50 from the binding curve, but the shorter 1-46 construct produced greater stabilization (Fig. 1E, dotted lines; Tm50 ∼0.5 °C higher than full-length), consistent with the ITC data. Addition of capsid hexamers produced minor changes in the PQBP1 far-UV circular dichroism (CD) spectrum (200-250 nm), reflecting altered secondary-structure content (Fig. S1C). These changes required an intact capsid R18 pore and were lost with the R18L mutant, indicating they arise from the N-terminal pore interaction (Fig. S1D). Van’t Hoff analysis of the CD thermal denaturation gave apparent unfolding enthalpies and melting temperatures for the WW-containing and full-length constructs (Fig. S1E). The 1-104 (Nt-WW) construct showed a higher apparent unfolding entropy than full-length. This suggests that a majority of PQBP1’s conformational entropy resides in the WW region. These observations are consistent with the WW domain driving the favorable binding entropy measured by ITC (Fig. 1C).

Together, the N-terminus drives high-affinity capsid binding, while regions through the WW domain shape the binding dynamics of the full-length protein. The favorable entropic contribution and length-dependent affinity suggest a binding-coupled conformational change within a disordered ensemble, which we next examined by NMR.

### Capsid binding perturbs conformational behavior within the WW domain

Given the central role of the WW domain in capsid hexamer binding and the extensive intrinsic disorder of full-length PQBP1 observed in prior NMR studies, we focused residue-level analyses on the Nt-WW construct (PQBP1 residues 1-104, Figs. S2A, S2B) (13, 16, 17, 18). Residue-specific responses to capsid binding were assessed by monitoring signal intensity and chemical shift changes by NMR spectroscopy during hexamer titration (Fig. 2A). Residues were classified as capsid-responsive when signal intensity decreased below 50% of the unbound value at the highest hexamer concentration tested (Fig. S2C). These experiments revealed pronounced line broadening and intensity loss for a subset of resonances upon titration (Figs. 2B, S2C), consistent with intermediate exchange and a binding-induced conformational change.

**Figure 2.**
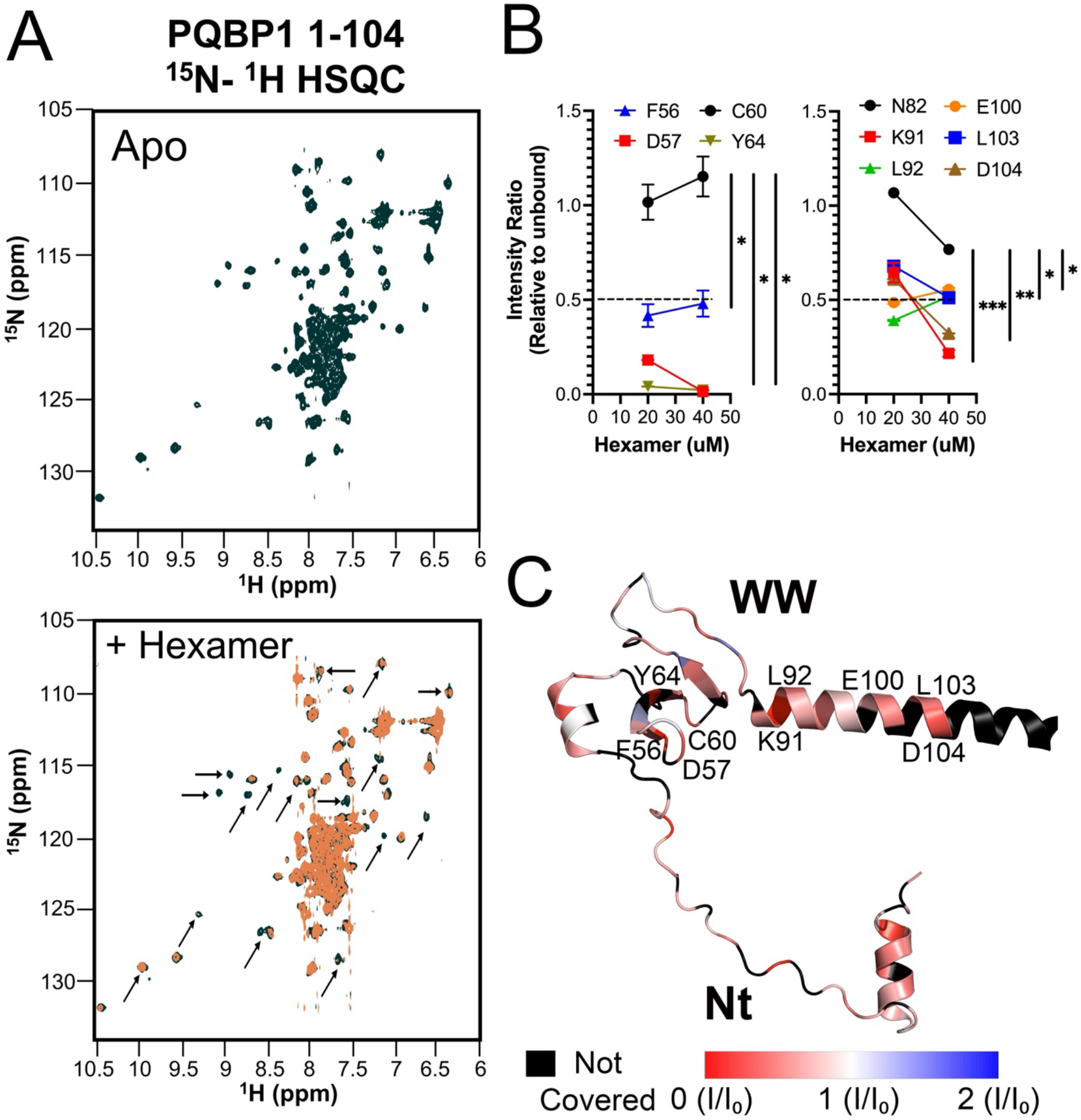
Capsid binding induces dynamics within the WW domain. (A) 2D ¹H-¹⁵N HSQC spectrum of ¹⁵N-labeled PQBP1 (residues 1-104) without (black) and with (orange) HIV-1 capsid hexamer (100 µM PQBP1, 40 µM hexamer). Selective peak broadening and intensity loss upon hexamer addition are indicated by arrows. Representative of three independent experiments. (B) Residue-specific intensity ratios (I/I₀) within the WW domain and adjacent linker, plotted as a function of increasing hexamer concentration. Residues with I/I₀ < 0.5 at 40 µM hexamer (dashed line) were considered capsid-responsive. At 40 µM hexamer, capsid-responsive residues were compared against neighboring non-responsive residues (C60 or N82) by unpaired t-test; *p<0.05, **p<0.01, ***p<0.001. (C) Hexamer-induced perturbations mapped onto a predicted PQBP1 (1-104) model, colored by intensity ratio (I/I₀) at 40 µM hexamer: unaffected residues (white), exchange-broadened (red), altered relaxation (blue). Perturbations concentrate in the WW domain and proximal linker. Model generated with AlphaLink2 using DSSO crosslinker-derived restraints for integrative modelling; visualized in PyMOL.

The most pronounced capsid-induced perturbations clustered within two regions of the WW domain. One set of effects was localized to the ∼50-64 segment, in which residues F56, D57, and Y64 exhibited enhanced line broadening upon titration. A second cluster was observed near the C-terminal end of the WW domain and the adjacent linker boundary (∼89-105), encompassing residues K91, L92, E100, L103, and D104 (Fig. 2B, dotted line). These residues were considered responsive relative to nearby unchanging residues (e.g., C60, N82), whose intensity ratios remained near 1. Mapped onto the Nt-WW AlphaLink2 model, these responsive residues concentrate in the WW domain and proximal linker, where intensity ratio is colored from red (exchange-broadened) through white (unaffected) to blue (increased intensity) (Fig. 2C).

Consistent with this WW-linker clustering, a nitroxide spin label (MTSL) introduced at C60, the sole cysteine in the WW domain, provided orthogonal confirmation by paramagnetic relaxation enhancement. This effect occurs when the unpaired electron of MTSL broadens NMR signals of nearby residues in a distance-dependent manner, chemical reduction of the label removes this effect to give a diamagnetic reference, and the ratio of the two (Iₚ/I_d_) reports proximity to the spin label over ∼10–25 Å (19, 20). Upon hexamer binding, the WW-linker residues L92, L103, and D104 showed increased PRE, indicating a sampling of conformations closer to C60 (Figs. S2D, S2E) and corroborating the titration-based broadening. This is consistent with the binding-coupled conformational shift predicted by the thermodynamics (Fig. 1).

### Hexamer binding promotes distal changes in PQBP1

Residue-level NMR resolved the Nt-WW response but could not report on the rest of full-length PQBP1. The size and disorder of the full-length protein led to severe crowding of overlapping disordered-residue signals, precluding reliable intensity measurements and backbone assignment (13, 16, 17, 18). To determine whether capsid engagement affects regions outside Nt-WW, we used hydrogen-deuterium exchange mass spectrometry (HDX-MS), which reports backbone solvent accessibility across the full-length protein. Regions that exchange deuterium more slowly upon binding are more shielded from solvent, indicating where capsid engagement alters local structure or contacts.

HDX-MS localized the capsid response to a broader region of the PRD (Fig. S3A). Within this region, peptide uptake decreased upon binding, and the peptide covering residues 180-196 showed a reproducible decrease across triplicate measurements (Fig. 3A). Reduced uptake here persisted across all timepoints from 30 seconds to 5 minutes, indicating a stable, prolonged change in solvent protection rather than a transient fluctuation. The Nt-WW region showed minimal deuterium uptake difference between bound and unbound states (Fig. S3B, peptides 9-26 and 46-72). Peptide-level HDX-MS may not be sensitive enough to capture the Nt-WW response. Since rapidly exchanging backbone amides in disordered regions saturate within the earliest timepoints, this can narrow the window for detecting protection on this timescale (21). Peptide averaging can also offset increases and decreases in exchange at adjacent residues (22). This fast-timescale behavior is instead resolved by NMR (Fig. 2).

**Figure 3.**
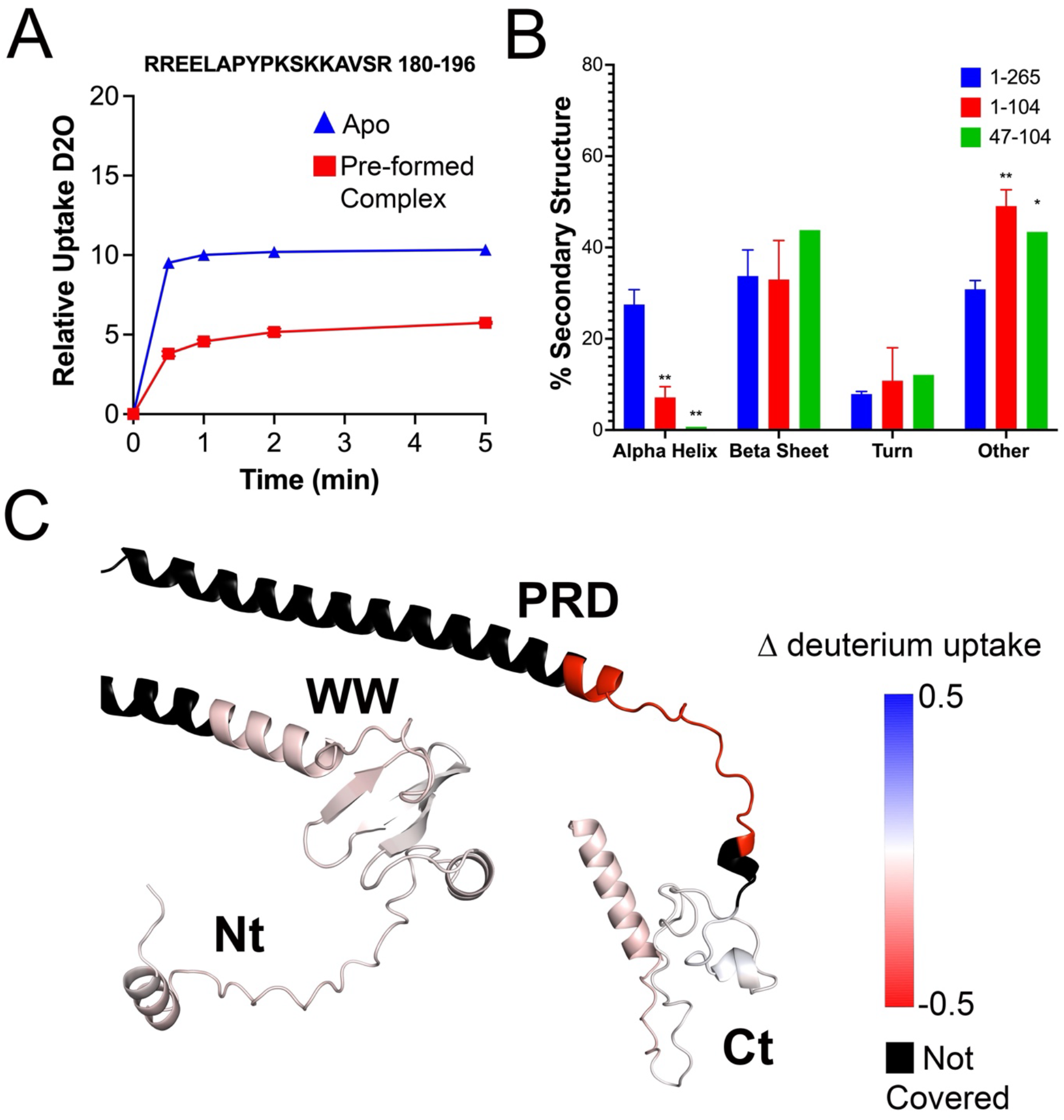
Hexamer binding promotes distal changes in PQBP1. (A) Hydrogen-deuterium exchange mass spectrometry (HDX-MS) representative relative deuterium uptake curves for PRD peptide (hexamer-bound vs. unbound). Biological triplicates are shown as mean ± SD (n=3). Significance assessed by DECA at 95% confidence intervals with back-exchange correction. Reduced uptake indicates decreased solvent accessibility in the distal PRD upon hexamer binding. Data is representative of three independent labeling experiments. (B) Far-UV CD spectra of PQBP1 FL and mutants at 25 °C under matched conditions. Secondary structure was estimated using BeStSel from biological replicates. Data represent the mean ± SD of two biological replicates. Statistics were determined by one-way ANOVA with Tukey’s post hoc test (*p < 0.05, **p < 0.01, ***p < 0.001, ****p < 0.0001). (C) Hydrogen-deuterium exchange mass spectrometry (HDX-MS) difference map (hexamer-bound minus unbound PQBP1) at 30 s (DynamiX, Waters). Decreased deuterium uptake (red) indicates hexamer-induced protection; increased uptake (blue) indicates increased dynamics. Sequence coverage: 66%; average peptide redundancy: 3.4. Analyzed using DECA with back-exchange correction. Data is representative of three independent labeling experiments (biological triplicate). Model generated with AlphaLink2 using DSSO crosslinker-derived restraints for modelling; visualized in PyMOL.

CD secondary-structure analysis of PQBP1 truncation constructs provides structural support for this interpretation. PQBP1 1-104 (Nt-WW) was predominantly β-rich, consistent with the reported triple-stranded β-sheet fold of the WW domain (13, 16), whereas full-length PQBP1 contained additional helical content absent from these truncations. Quantification confirmed reduced α-helical and increased disordered content in the 1-104 and 47-104 constructs relative to full-length (Fig. 3B). The AlphaLink2 model, constrained by experimental DSSO cross-link restraints (see Methods and Extended Data 2), represents an unbound conformation of PQBP1 (Fig. 3C) (32). In this model, the capsid-protected peptide (residues 180-196) maps to a loop extending from an α-helix in the PRD (Fig. 3C). Flexible loops can act as hinges, allowing the sampling of multiple conformations (33). Together, these observations suggest that hexamer binding either promotes ordering of this loop or shifts PQBP1’s conformational equilibrium toward a more compact state, reducing solvent accessibility within the PRD.

Together, these data show that hexamer binding at the N-terminus produces a stable reduction in solvent accessibility within the distal PRD, extending the capsid response beyond the binding surface. This positions the PRD, rather than the N-terminal contact alone, as a candidate mediator of the sensing events downstream of capsid recognition.

### WW-linker and PRD regions are required for infection-dependent cGAS recruitment

Prior work established that PQBP1 recruits cGAS through regions distinct from its N-terminal capsid-binding surface, and that this recruitment requires PQBP1 to interact with the capsid (8, 9). Our NMR and HDX data show that capsid binding at the N-terminus reorganizes distal regions of PQBP1, the WW-linker, and the PRD (Figs. 2, 3). We therefore reasoned that these distal regions, though outside the capsid-binding interface, might be the elements that couple capsid engagement to cGAS recruitment. To test this, we used loss-of-function mutagenesis to distinguish the capsid-binding surface from the capsid-responsive regions and determine which is required for cGAS recruitment in infected cells.

Guided by the mapped capsid-responsive residues (Figs. 2, 3), we introduced alanine substitutions across the capsid-responsive regions: the WW-linker (89-93Ala and 100-105Ala) and the distal PRD (189-192Ala) (Fig. 4A). These residues lie outside the primary N-terminal hexamer-binding site (residues 1-46), so their mutation uncouples the direct capsid contact from the distal conformational response. WT and mutant PQBP1-sfYFP constructs were stably transduced into THP-1 Flag-cGAS cells, and PQBP1-sfYFP and Flag-cGAS expression was confirmed by sorting, fluorescence, and immunoblotting (Figs. S4A, S4B). All PQBP1 constructs were expressed at comparable levels, so the infection-dependent differences below are not attributable to construct abundance.

**Figure 4:**
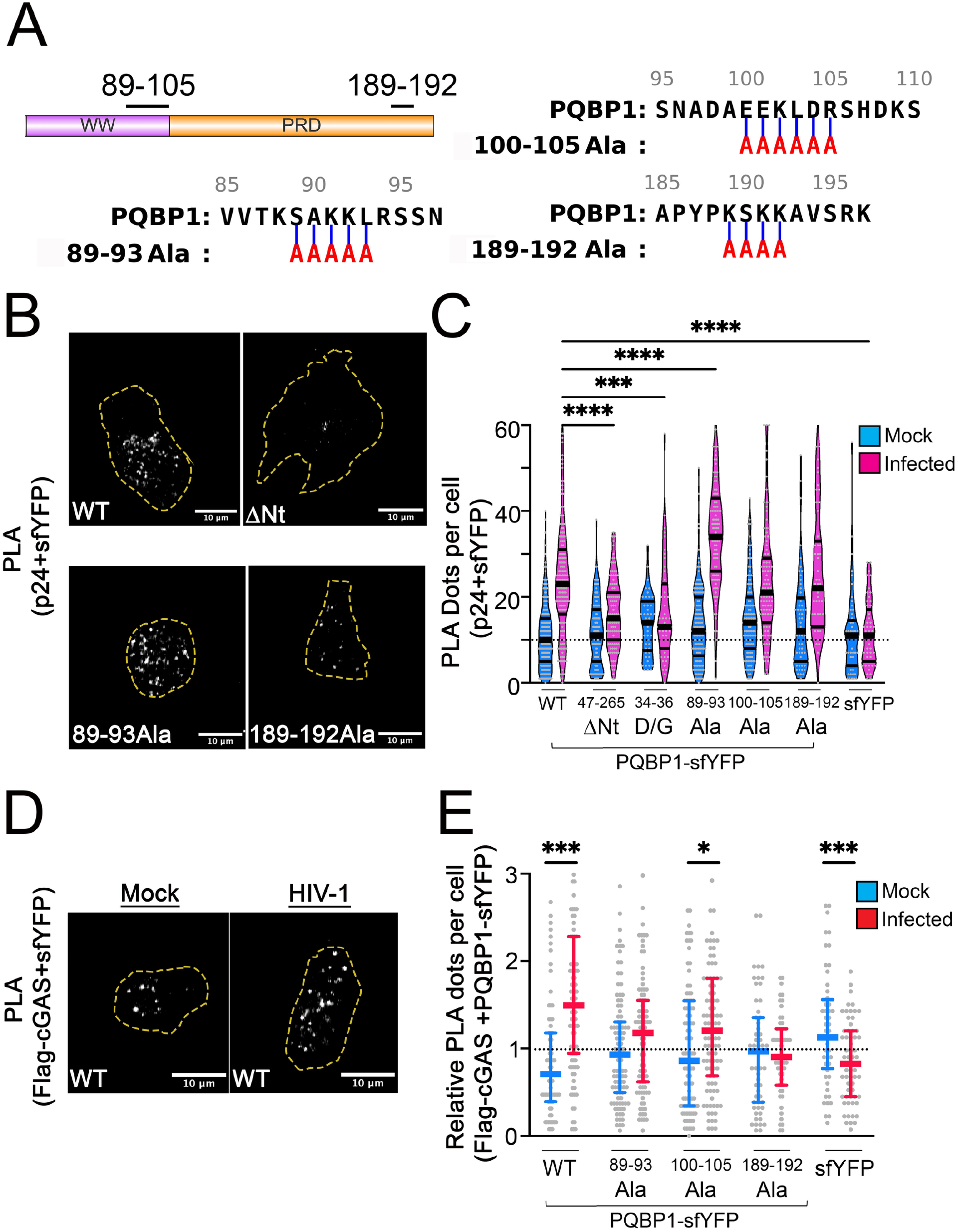
Residues within capsid-responsive regions contribute to cGAS recruitment. (A) PQBP1 domain schematic (WW, PRD) indicating alanine substitutions (red) in CA hexamer binding-responsive regions identified by NMR and HDX-MS: WW-linker (89-93, 100-105), and distal PRD (189-192). (B) Representative confocal images of p24-PQBP1 proximity ligation assay (PLA) in HIV-1-infected THP-1 cells stably expressing the indicated PQBP1-sfYFP construct [WT, ΔNt (47-265), D/G (D34G, D35G, D36G), 89-93Ala, 100-105Ala, and 189-192Ala]. ΔNt and D/G are capsid binding-deficient mutants and serve as negative controls. PLA detects PQBP1-sfYFP and HIV-1 capsid (p24) proximity (<50 nm). Scale bar, 10 µm; dashed outline indicates the cell boundary. (C) Quantification of p24-PQBP1 PLA puncta per cell across constructs in mock (blue) and HIV-1-infected (magenta) cells. Bars show median and 10-90% interquartile range. Because no capsid antigen is present in mock conditions, infected cells were compared across constructs by one-way ANOVA with Šídák’s post hoc test relative to wild type (WT) PQBP1. ***p < 0.001, ****p < 0.0001. Representative of at least two independent experiments. (D) Representative confocal images of proximity ligation assay (PLA) signal in THP-1 cells stably expressing PQBP1-sfYFP and Flag-cGAS under mock or HIV-1/Vpx infection. PLA detects PQBP1-sfYFP and Flag-cGAS proximity (<50 nm). Scale bar: 10 µm; dashed outline indicates cell boundary. (E) Quantification of cGAS-PQBP1 PLA puncta per cell, expressed relative to the median of the mock-infected condition for each construct, in mock (blue) and HIV-1-infected (red) cells. Bars show median and 10-90% interquartile range. The effects of infection and construct were assessed by two-way ANOVA; infected and mock conditions were compared within each construct by Šídák’s post hoc test. *p < 0.05, ***p < 0.001. Pooled single cells from three independent experiments

To measure capsid interaction in cells, we used an established proximity ligation assay (PLA) between the incoming capsid (HIV-1 p24) and PQBP1-YFP, visualized by immunostaining and confocal microscopy. At 2 hours post-infection, we quantified the number of PLA puncta per cell and compared the median values (Figs. 4B, 4C) (9). The sfYFP-only expression and mock infection showed few puncta, establishing that the signal is specific to both exogenously expressed PQBP1 and HIV-1 infection (Fig. 4C). The capsid-binding-defective ΔNt (a.a. 47-265) and D/G (D32G, D33G, D34G) PQBP1 constructs (11) gave near-background signal, defining the lower bound of the assay (Figs. 4B, 4C). Against this window, HIV-1-infected cells expressing wild-type PQBP1 showed abundant capsid-proximal puncta. Both the WW-linker mutants (89-93Ala and 100-105Ala) and the PRD mutant (189-192Ala) PQBP1s retained puncta at or above full-length levels (Figs. 4B, 4C), indicating the WW-linker and PRD substitutions do not disrupt the primary capsid interaction.

Because these mutants retained capsid binding, we next evaluated cGAS recruitment efficiency of the mutants by PLA between PQBP1-sfYFP and Flag-cGAS. PQBP1-cGAS puncta frequencies were near background in uninfected cells, and HIV-1 infection increased the puncta frequency, shown by ∼ 2-fold increase in the median value compared to mock-infected, establishing infection-dependent PQBP1 and cGAS association (Figs. 4D, 4E, S4C) (8, 9). Both the wildtype and the 100-105Ala mutant showed an infection-dependent increase, whereas the 89-93Ala and 189-192Ala mutants did not (Fig. 4E). Because 89-93Ala and 189-192Ala retain capsid binding, yet fail to recruit cGAS, these regions are required for cGAS recruitment post to capsid engagement (Fig. 4C).

Together, these results show that the WW-linker (89–93) and PRD (189-192) regions are structurally distinct from the primary capsid interface yet functionally required for cGAS association. These findings indicate that capsid-induced conformational changes in these regions are necessary for productive complex formation for sensing of HIV-1 infection.

## DISCUSSION

cGAS detects cytosolic DNA to initiate type I interferon responses, yet HIV-1 presents an unusual challenge for this pathway. During infection, reverse-transcribed viral DNA is low in abundance, transiently exposed, and shielded within the viral capsid, limiting its accessibility to cGAS. Our findings lend further support to a model in which PQBP1 overcomes this challenge through a two-step mechanism: PQBP1 first recognizes the HIV-1 capsid and subsequently recruits cGAS to the viral replication complex. We propose that capsid engagement is accompanied by changes in PQBP1 conformational organization that promotes cGAS recruitment. Capsid binding influences conformational behavior within both the WW-linker and distal PRD, and PLA demonstrates that these regions are required for infection-dependent co-localization of PQBP1 and cGAS. NMR localizes the responsive residues to the WW-linker, whereas HDX-MS identifies protection within the PRD. Although each measurement provides an indirect view of conformational behavior, together with the cellular data, they support a model in which capsid engagement is associated with structural changes in PQBP1 that accompany productive cGAS recruitment. The thermodynamics of this interaction reinforce this interpretation: apparent affinity decreases with increasing construct length, and binding is entropically favorable across constructs, consistent with distal regions influencing the conformational ensemble sampled during capsid binding (15, 23).

The WW-linker and PRD lie outside the N-terminal R18 pore-binding region, yet both are required for productive cGAS recruitment. The WW-linker contains residues identified by NMR as capsid responsive, whereas the PRD encompasses the region protected by HDX-MS following capsid engagement. Together, these observations identify two functionally important regions outside the primary capsid-binding interface. The structural basis of these changes remains incompletely resolved. NMR identifies capsid-responsive residues within the WW-linker, whereas HDX-MS demonstrates altered solvent accessibility within the distal PRD. Together, these complementary approaches indicate that capsid engagement influences regions well beyond the primary capsid-binding interface, although the precise structural rearrangements remain to be defined.

The functional importance of the WW domain and distal regions of PQBP1 is further supported by prior studies. The Renpenning syndrome-associated Y65C mutation disrupts WW-domain folding and impairs PQBP1-dependent innate immune signaling, including cGAS recruitment (8, 16). Likewise, patient-derived mutations truncating the PRD and C-terminal regions compromise HIV-1 innate sensing (8). Although these variants do not map directly to the residues examined here, together they support the conclusion that both WW-domain integrity and distal regions of PQBP1 are required for productive cGAS recruitment.

This behavior reflects dynamic coupling, in which a disordered protein uses ligand-induced conformational changes to couple recognition to downstream function (15, 23). We propose that capsid engagement remodels the PQBP1 conformational ensemble to promote cGAS recruitment to the viral reverse transcription complex. This model provides a potential solution to a central challenge in HIV-1 innate sensing: reverse-transcribed viral DNA is low in abundance, transiently exposed, and largely shielded within the viral capsid. By first recognizing the capsid, PQBP1 recruits and locally enriches cGAS at the site of viral DNA synthesis before sufficient DNA is exposed for activation. Our biophysical measurements used isolated hexamers, which reproduce the pore interaction but not the curvature or avidity of an assembled core; because the lattice presents multiple PQBP1-binding sites, multivalent engagement could further increase the local abundance of PQBP1, resulting in enhanced cGAS recruitment to the reverse transcription complex, potentially increasing sensitivity to scarce viral DNA beyond what is captured by single-hexamer measurements. How avidity within the assembled capsid lattice shapes this response, whether PQBP1 also modulates cGAS catalytic activity, and whether it influences the threshold of viral DNA required for cGAS activation remain important questions for future study.

Together, these results indicate that capsid binding and cGAS recruitment depend on distinct regions of PQBP1 and support a model in which capsid recognition reorganizes PQBP1 conformational behavior to promote cGAS recruitment to the viral reverse transcription complex. This mechanism may extend beyond HIV-1 to other PQBP1 ligands, including oligomeric assemblies such as tau aggregates (10, 24), suggesting that ligand-induced conformational remodeling may represent a broader mechanism by which PQBP1 couples recognition of multivalent ligands to selective innate immune activation (25).

## METHODS

### Protein expression and purification

Human PQBP1 constructs (full-length 1-265, N-terminus 1-46, and domain truncations) were cloned into bacterial expression vectors containing N-terminal His6-TEV, His6-or C-terminal His6-tags or purchased from Lifetein LLC with >90% purity. His-tagged constructs were expressed in E. coli BL21(DE3) cells grown in LB or M9 minimal media, as appropriate. Cultures were induced with 0.5 mM IPTG at OD600 ∼0.6 and incubated overnight at 20 °C. Cells were harvested by centrifugation and lysed by sonication in lysis buffer containing 50 mM NaH₂PO₄, 300 mM NaCl, 10 mM imidazole, pH 8.0. Clarified lysates were applied to HisTrap FF Ni-NTA resin (Cytiva), washed extensively, and eluted with 250 mM imidazole. Where applicable, tags were removed by overnight TEV protease digestion at 4 °C, followed by reverse Ni-NTA purification. Final purification was performed by either HiTrap anion exchange chromatography or size-exclusion chromatography using a Sephacryl 26/60 column equilibrated in 20 mM HEPES, 100 mM KCl, pH 8.0.

HIV-1 capsid hexamers were expressed and purified as previously described, using disulfide-stabilized hexamer constructs. Hexamers were dialyzed into matching buffer prior to all experiments (11, 26). Biophysical experiments were performed in buffer conditions optimized for each technique. CD and NMR measurements were conducted in a sodium phosphate buffer (20 mM, pH 7.2) to minimize contributions from far-UV absorbance. ITC was performed in HEPES buffer (20 mM HEPES, 100 mM KCl, 2.5 mM MgCl₂, pH 8.0) to avoid phosphate interference with binding thermodynamics. HDX-MS was performed in HEPES buffer (pH 8.0) compatible with inline pepsin digestion. All buffers were matched for ionic strength and pH within each experiment. The consistency of thermodynamic parameters across ITC and CD-monitored thermal denaturation (van’t Hoff analysis) performed in independent buffer systems supports the robustness of the observed binding signatures.

### Isothermal titration calorimetry

Isothermal titration calorimetry (ITC) experiments were performed using a MicroCal PEAQ-ITC instrument at 25 °C. PQBP1 proteins were dialyzed extensively against assay buffer (20 mM HEPES, 100 mM KCl, 2.5 mM MgCl₂, pH 8.0). Capsid hexamers were loaded into the syringe and titrated into the sample cell containing PQBP1. Typical experiments consisted of an initial 0.4 μL injection, followed by 18-20 2 μL injections with a 150 s spacing. Data were integrated and fit using a single-site binding model in the manufacturer’s software. Reported thermodynamic parameters represent mean values from at least two independent experiments. The equilibrium dissociation constant (K_d_) and binding enthalpy (ΔH) were obtained directly from the fits, and the free energy change (ΔG) and entropic contribution (TΔS) were calculated using standard thermodynamic relationships (ΔG = RT ln K_d_; TΔS = ΔH − ΔG).

### Nano-differential scanning fluorometry

Differential scanning fluorimetry was performed using a NanoTemper Prometheus NT.48 instrument. Stabilized HIV-1 capsid hexamers were prepared at a final concentration of 2 µM in assay buffer (20 mM Tris-HCl, 50 mM NaCl, pH 7.5). PQBP1 or PQBP1 truncation constructs were titrated into the hexamer solution at the indicated concentrations and equilibrated for 10 min at room temperature prior to analysis. Samples were loaded into standard capillaries and heated from 20 °C to 95 °C at a ramp rate of 1 °C min⁻¹. Thermal unfolding was monitored by intrinsic tryptophan fluorescence at 330 nm and 350 nm, and melting curves were calculated from the fluorescence intensity ratio (350/330). Apparent melting temperatures (T_m)_ were determined from the first derivative of the fluorescence ratio using NanoTemper PR ThermControl software. All measurements were performed in at least duplicate, and representative melting profiles are shown.

### Circular dichroism spectroscopy

Circular dichroism (CD) spectra were collected on a Jasco J-1500 spectropolarimeter using quartz cuvettes with a 1-mm path length. Spectra were recorded at 20 °C, with a 1-nm bandwidth and a scan rate of 50 nm min⁻¹. Protein concentrations were determined by A280 and were typically 8 µM for PQBP1 constructs (0.24 mg mL⁻¹). Capsid assemblies were prepared at 8 µM CA (p24-equivalent; 0.19 mg mL⁻¹), corresponding to 1.33 µM hexamer. For binding experiments, PQBP1 (8 µM) was mixed with capsid at 1.33 µM CA-equivalent, yielding a total protein concentration of approximately 0.43 mg mL⁻¹. Where indicated, component-subtracted spectra were generated using matched buffer baselines, with spectra from hexamer-only controls subtracted from PQBP1-capsid mixtures to isolate ligand-dependent changes near 225 nm. Spectra were processed with CDXtools and plotted with GraphPad Prism 9. PQBP1 mutant proteins were similarly analyzed at 8 µM. Secondary structure content was estimated from the CD spectra using the BeStSel algorithm.

For thermal denaturation experiments, ellipticity at 225 nm was monitored as a function of temperature, and unfolding curves were analyzed using a published monomeric two-state integrated van’t Hoff model (27) with linear native and unfolded baselines. The unfolding equilibrium constant was expressed using the integrated van’t Hoff equation:

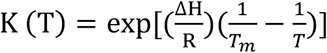

where ΔH is the unfolding enthalpy (assumed temperature-independent, ΔCp = 0), Tm is the melting temperature, R is the gas constant, and T is the absolute temperature. The fraction unfolded was related to the equilibrium constant by:

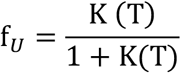

Thermodynamic parameters (ΔG, ΔH, and TΔS) were obtained by nonlinear least-squares fitting of the unfolding transitions under the assumption of a reversible two-state process.

### Crosslinking Mass Spectrometry (XL-MS) constrained Alphalink2 model

Recombinant PQBP1 and stabilized HIV-1 capsid hexamers (30 µM each) were crosslinked with DSSO (1:40 molar ratio, 30 min, room temperature) in 20 mM HEPES, 100 mM NaCl, pH 7.5. Reactions were quenched with 75 mM Tris, and complexes purified by Ni-NTA and buffer-exchanged. Samples were denatured, reduced, and alkylated (8 M urea, 10 mM TCEP, 40 mM CAA), digested with 0.2 µg Trypsin/Lys-C (Promega) overnight (30 °C). Following digestion, samples were acidified with TFA, and approximately 100 ng of peptides were loaded onto EvoTips (EvoSep) according to the manufacturer’s instructions. Peptide separation was performed on an EvoSep One system (EvoSep) coupled to a timsTOF Pro 2 mass spectrometer (Bruker Daltonics) using a 15-cm Aurora Elite CSI column (IonOpticks) and the 20 SPD Whisper Zoom gradient. Mass spectrometry data were acquired in positive ion mode DDA-PASEF. The ion mobility acquisition time was set to 200 ms, with 6 MS/MS scans per cycle. A minimum precursor intensity threshold of 3,000 and a target intensity of 50,000 were applied. Only precursor ions with charge states between 3 and 8 were selected for fragmentation. Collision energy was applied linearly as a function of ion mobility, ranging from 0.73 at 23 eV to 1.63 at 95 eV. Raw data were converted to MGF (MSConvert, v3.0.24336) and searched with XiSearch/XiFDR (v1.8.7) against a custom database containing the purified proteins of interest (20 ppm tolerance) (28, 29). DSSO was specified to react with lysine, serine, threonine, tyrosine, and protein N-termini, consistent with its established primary amine and secondary hydroxyl reactivity. Carbamidomethylation (Cys) was set as a fixed modification; methionine oxidation, N-terminal acetylation, and DSSO mono-links/hydrolyzed and amidated crosslinker stubs were variable. ^14^N/^15^N labeling was included (29, 30, 31). The dataset was filtered at a 5% false-discovery rate (FDR). Individual crosslinks were additionally evaluated by the number of supporting peptide-spectrum matches (PSMs) and reproducibility across the ¹⁴N/¹⁵N-labeled samples. Inter and intra-protein crosslinks supported by multiple PSMs were retained for interface assignment, whereas single-PSM or sequence-ambiguous crosslinks were treated as low-confidence and excluded. DSSO-derived restraints were incorporated into AlphaLink2 modeling for unbound PQBP1 following default parameters, and crosslinked residues are reported in Extended Data 2 (32). The resulting model was refined using GalaxyWeb.

### NMR spectroscopy

Uniformly ¹⁵N-labeled PQBP1 (residues 1-104) was expressed in *E. coli* grown in M9 minimal medium supplemented with ¹⁵N ammonium sulfate as the sole nitrogen source and ¹³C glucose as the sole carbon source. NMR experiments were performed at 298 K on a Bruker Neo 800 MHz spectrometer equipped with a 5 mm TCI z-axis gradient cryogenic probe. Samples were prepared in 20 mM sodium phosphate, 50 mM NaCl, 1 mM DTT, 10% D₂O, pH 7.2. Two-dimensional ¹H-¹⁵N HSQC spectra were acquired for 100 µM PQBP1 alone and following titration with increasing concentrations of HIV-1 capsid hexamers. Spectra were processed in TopSpin and analyzed in NMRFx Analyst. Peak lists and intensity analyses were generated in NMRtist, based on assignments obtained with ARTINA analysis of ¹⁵N and ¹³C 2D planes (Extended Data 1) (34, 35). Per-residue assignment confidence is reported in Supplementary Figure 2. Residues were classified as capsid-responsive by three criteria: (i) a reduction in backbone amide peak intensity below 50% of the unbound value (I/I₀ < 0.5) at the highest hexamer concentration tested; (ii) a monotonic, hexamer-concentration-dependent decrease in intensity across the titration; and (iii) significant attenuation relative to adjacent non-responsive residues at the highest hexamer concentration, by unpaired t-test. Residues meeting all three criteria were considered to undergo capsid-induced line broadening consistent with intermediate exchange.

PQBP1 1-104 was spin-labeled at C60, the only cysteine in the construct, by overnight incubation with a 10-fold molar excess of (1-oxyl-2,2,5,5-tetramethyl-Δ3-pyrroline-3-methyl) methanethiosulfonate (MTSL; Sigma-Aldrich) at 4 °C in the dark. Complete conjugation was confirmed by intact-protein MALDI mass spectrometry, and excess MTSL was removed by dialysis against 20 mM sodium phosphate, 50 mM NaCl, pH 7.2. 2D ¹H-¹⁵N HSQC spectra were acquired at 100 µM ¹⁵N-PQBP1 on the Bruker 800 MHz spectrometer using the parameters above. Paramagnetic spectra were collected first, followed by diamagnetic controls after addition of 2 mM sodium ascorbate.

Peak intensities were extracted in NMRFx Analyst at curated backbone amide assignments and computed per replicate (n = 2) as the paramagnetic-to-diamagnetic ratio (Iₚ/I_d)_, where lower values indicate closer proximity to C60. Ratios with diamagnetic intensity below 3× spectral noise were excluded. Labeling was confirmed by the C60-proximal residues D57 and Y64, fully broadened in the apo paramagnetic spectrum (Iₚ/I_d_ ≈ 0) and restored in the reduced diamagnetic reference, confirming that intensity loss was paramagnetic in origin. The WW-linker residues showed a range of apo Iₚ/I_d_ values reflecting their differing proximity to C60 (K90 = 0.73 ± 0.09, L103 = 0.70 ± 0.21, D104 = 0.55 ± 0.03, L92 = 1.25 ± 0.30). Upon hexamer binding, L92, L103, and D104 decreased (L92, 1.25 ± 0.30 to 0.17 ± 0.11; L103, 0.70 ± 0.21 to 0.09 ± 0.04; D104, 0.55 ± 0.03 to 0.22 ± 0.04), indicating increased proximity to C60. The K90 amide was not resolved in the bound state and was not determined. Distances can be estimated following the Battiste and Wagner approach (19, 20): Iₚ/I_d_ < 0.15 corresponds to ∼12 Å from the spin label, 0.15–0.50 to ∼12–20 Å, and 0.50–0.85 to ∼20–25 Å (∼8 Å accounts for the MTSL linker). Bound-state ratios were assessed relative to each residue’s apo value rather than as absolute distances, since the increased rotational correlation time of the complex can broaden resonances globally.

### Hydrogen-deuterium exchange mass spectrometry

HDX-MS experiments were performed at the UC San Diego HDX-MS Core Facility using a Waters HDX platform coupled to a SYNAPT® G2-Si mass spectrometer. 6 µM of PQBP1 samples with or without 3 µM HIV-1 hexamers were equilibrated at 25 °C for 10 min. Occupancy was estimated using the quadratic binding equation accounting for ligand depletion at the experimental concentrations (6 µM PQBP1, 3 µM hexamer, Kd 6.2 µM), yielding ∼25% bound PQBP1 under these conditions. Deuterium exchange was initiated by a 16-fold dilution into deuterated buffer, yielding a final 93.75% D₂O concentration. Exchange reactions were carried out at 25 °C and sampled at 30 s, 1 min, 2 min, and 5 min. Samples were subjected to inline acidic quenching followed immediately by online pepsin digestion and low-temperature chromatographic separation prior to mass spectrometric analysis. Undeuterated control peptides were identified using ProteinLynx Global SERVER (PLGS, Waters). DynamiX (Waters) was used for peptide identification, calculation of deuterated peptide uptake, and data quality control. DECA was used for downstream HDX analysis, statistical evaluation, and data visualization. Back-exchange-corrected deuterium uptake values, generated in DECA, were used for structure-based PDB mapping, with relative fractional uptake differences averaged over the 0 to 5 min time window. Raw (uncorrected) uptake values were used for butterfly plots and relative uptake plots, consistent with qualitative comparisons of exchange behavior. Statistical significance was assessed using 95% confidence limits derived from DECA. Data available in Extended data 3.

### Mutagenesis

PQBP1-mVenus (sfYFP) constructs were generated by NEBuilder HiFi DNA Assembly (NEB). The insert was amplified by PCR with Q5 High-Fidelity 2X Master Mix (NEB) using the mVenus primers below, and the vector was prepared by digesting pLenti6-PQBP1-eYFP with BamHI-HF and BsrGI-HF (NEB). Inserts and vectors were purified (NucleoSpin Gel and PCR Clean-up, Macherey-Nagel) and assembled by NEBuilder HiFi DNA Assembly. The sfYFP-only control vector was generated using primers that delete the PQBP1 coding sequence, leaving mVenus (sfYFP) alone.

Point mutants and truncations of PQBP1 were generated as previously described using Q5 site-directed mutagenesis (NEB) or commercial gene synthesis (SynBio) and verified by Sanger sequencing (9). Primer sequences are listed below.

mVenus-F: GACGGTACCGCGGGCCCGGGATCCACCGGTCGCCACCATGGTG

mVenus-R: CCCTCTAGACTCGATTACTTGTACAGCTCGTCCATGCCGAGAGTG

ΔPQBP1-sfYFP-F: AGCTTATGAGAATTCTGCAGTCGACGGTACC

ΔPQBP1-sfYFP-R: GAATTCTCATAAGCTTGAGCTCGAGATCTGAG

89-93Ala-F: GCGGCGTCATCCCATGCAGATG

89-93Ala-R: CGCCGCGGCCGATTTGGTAAC

100-105Ala-F: GCGGCGGCGAGCCATGACAAGTCG

100-105Ala-R: CGCCGCCGCAGCATCTGCATGGGATG

189-192Ala-F: CGGCGGCGGCAGTAAGCCGAAAG

189-192Ala-R: CCGCCGCATAGGGAGCCAGCTC

### Proximity Ligation Assay (PLA) and THP1 infection

For THP-1 experiments, cells were transduced with lentiviral constructs encoding PQBP1-sfYFP (full-length or mutant) and Flag-cGAS, as previously described (9). Double-positive populations were isolated by fluorescence-activated cell sorting (Bigfoot), and comparable PQBP1-sfYFP and Flag-cGAS expression across lines was confirmed by sorting, mean fluorescence intensity, and immunoblot (Figs. S4A, B). Sorted lines were differentiated with PMA, transfected with siRNA against endogenous PQBP1, and seeded at 2.0 × 10⁴ cells per well as previously described (9). Cells were fixed 2 h post spin-infection, permeabilized in 0.2% Triton X-100, blocked, and incubated overnight with rabbit anti-GFP (1:1000; Takara 632592) and mouse anti-Flag M2 (1:400; Sigma F1804) to detect the PQBP1-sfYFP-cGAS interaction, or with rabbit anti-GFP and mouse anti-p24 (HIV-1 capsid; ab9071) to detect the PQBP1-sfYFP-capsid interaction. Proximity ligation was performed using the Duolink In Situ Far-Red detection kit (Sigma) according to the manufacturer’s instructions, generating a punctate signal only when the two epitopes were within <50 nm. PLA puncta were imaged on a Zeiss LSM880 confocal microscope with a Plan-Apochromat 40×/1.4 Oil DIC or 63×/1.4 Oil DIC (Carl Zeiss Microscopy). Reactions containing only one primary antibody (anti-GFP alone, anti-Flag alone, or anti-p24 alone), and reactions substituting matched isotype IgG for a primary antibody, produced no PLA signal above background after CYTELY gating and analysis. Cell identification and per-cell puncta quantification were performed using CYTELY AI analysis (36); equivalent results were obtained using ImageJ Analyze Particles, as previously described (9).

VSVG-pseudotype HIV-1 and VLP-Vpx were generated and used as previously described (8, 9). Following siRNA transfection, PMA-differentiated THP1 cells were infected with a VSVG-pseudotype HIV-1 in the presence of VLP-Vpx. Approximately 5 ng of p24 or 2.5-5 RT units of HIV-1 were used per 2.0 x 10^4^ PMA-THP1 cells in 100 µL of media. Productive infection was confirmed by Renilla-Luciferase detection (Bright-Glo, PerkinElmer).

## STATISTICAL ANALYSIS

All quantitative data are presented as mean ± SD unless otherwise stated. For the p24-PQBP1 PLA, in which no capsid antigen is present in mock conditions, infected cells were compared across constructs by one-way ANOVA with Šídák’s post hoc test relative to full-length PQBP1. For the cGAS-PQBP1 PLA, in which both proteins are co-expressed, proximity signal was analyzed by two-way ANOVA to assess the effects of HIV-1 infection and PQBP1 construct. Following confirmation of a significant infection effect, infected and mock conditions were compared within each construct using Šídák’s post hoc test. Circular dichroism comparisons between constructs were performed by one-way ANOVA with Tukey’s post hoc test relative to full-length. All analyses were performed in GraphPad Prism, except HDX-MS, which was evaluated in DECA (95% confidence limits).

## DATA AVAILABILITY STATEMENT

The chemical crosslinking mass spectrometry data have been deposited to the ProteomeXchange Consortium via the PRIDE partner repository under accession PXD080336. The hydrogen-deuterium exchange mass spectrometry data have been deposited to MassIVE under accession MSV000102525. All reviewer credentials will be removed and the datasets released upon publication.

## Supporting information

Data S2. DSSO cross-linking mass spectrometry data for PQBP1.

Data S1. NMR chemical shift perturbation data for PQBP1.

## ACKNOWLEDGMENTS

We thank members of the Chanda laboratory for helpful discussions and feedback throughout this work. We are grateful to the Scripps Research core facilities for technical support, including Phil Ordoukhanian and the Biophysics and Biochemistry (B2) Core, the Microscopy Core (Scott Henderson and Kathryn Spencer), the Protein Production and Antibody Core, the NMR Core (Gerard J. Kroon), and the Mass Spectrometry Core. We thank Peter Wright and Maria Martinez-Yamout for assistance with NMR experimental design, analysis, troubleshooting, and review. We thank Steve Silletti (Biomolecular and Proteomics Mass Spectrometry Facility, University of California, San Diego) for HDX-MS assistance with data acquisition, peptide assignment, and back-exchange analysis. D.S.A. thanks Sergio Catz and Jennifer Johnson for their training and mentorship during the early stages of his scientific development at Scripps Research. We are grateful to Renate König and members of her laboratory, including Domenico Rizzo, for collaborative discussions and support. We thank João Mamede for providing PQBP1 plasmids. We also thank the members of the dissertation committee, Jeffrey W. Kelly, Michael Bollong, Andrew Ward, and Ashok Deniz, for valuable guidance and scientific insight. D.S.A. was supported by an NIH F31 award (NINDS 1F31NS141616-01). C.M.B. was supported by a Swedish Research Council grant (2023-00510).

## CONTRIBUTIONS

Conceptualization: D.S.A., S.M.Y., S.K.C., O.P. Methodology: D.S.A., A.M., T.Z., C.M.B., B.K.G.-P., O.P. Validation: D.S.A., A.M., S.M.Y. Formal Analysis: D.S.A., C.M.B., A.M. Investigation: D.S.A., A.M., T.Z., C.M.B., S.O., S.P., A.Me., D.G. Resources: B.K.G.-P., O.P. Visualization: D.S.A., A.M., S.M.Y. Writing - Original Draft: D.S.A. Writing - Review & Editing: D.S.A., S.M.Y., S.K.C., O.P., B.K.G.-P., A.M., J.P. Supervision: S.M.Y., S.K.C., O.P. Project Administration: D.S.A., S.M.Y., S.K.C. Funding Acquisition: S.M.Y., S.K.C.

## FUNDING

This study was funded by NIH R01-AI127302 R01-AI177265, NIH U54-AI170856. The Synapt HDX-MS instrument at BPMSF is supported by NIH shared instrumentation grant S10 OD016234.

## AI STATEMENT

Manuscript preparation included the use of Claude (Anthropic) for editorial assistance with clarity and writing. Data shared during this process is covered under Scripps Research institutional licensing terms and has not been used for model training.

---

**Supplemental Figure 1:**
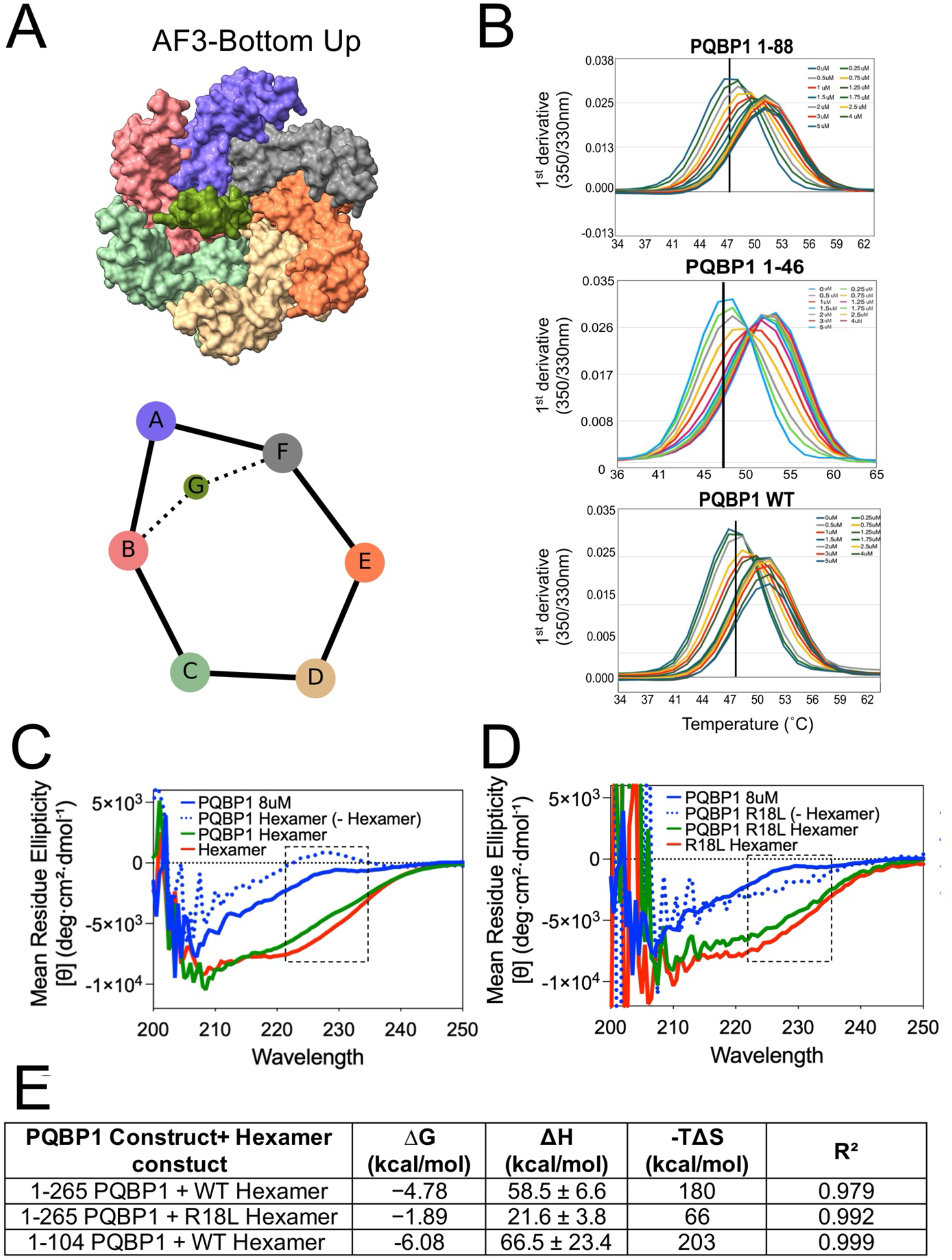
Regions outside Nt regulate hexamer binding. (A) Structural model of the PQBP1 N-terminus (Nt, green) engaging the HIV-1 capsid hexamer (subunits A-F). Top, ChimeraX visualization of an AlphaFold3-derived Nt-bound model docked onto the hexamer surface. Bottom, schematic contact diagram showing the relative positioning of PQBP1 interaction sites across adjacent capsid subunits. (B) First-derivative nanoDSF profiles of HIV-1 capsid hexamer in response to the increasing concentration of indicated PQBP1 protein, plotted against temperature. Shifts in melting temperature (Tm) reflect construct-dependent stabilization of the hexamer. Representative of at least two independent experiments. (C) Far-UV circular dichroism (CD) spectra of PQBP1 FL with and without WT capsid hexamer under matched conditions, shown as mean residue ellipticity (MRE) across 200-260 nm. Differences in ellipticity reflect hexamer-dependent changes in secondary structure. Representative of three independent experiments. (D) Far-UV CD spectra of PQBP1 FL with and without the binding-deficient R18L capsid hexamer. Representative of two independent experiments. (E) Thermal denaturation monitored by CD at 225 nm. Representative melting curves for PQBP1 alone and with hexamer, corresponding to the conditions in C and D. Right, thermodynamic parameters of the unfolding transitions derived from replicate measurements using an integrated two-state van’t Hoff model (see Methods).

**Supplemental Figure 2:**
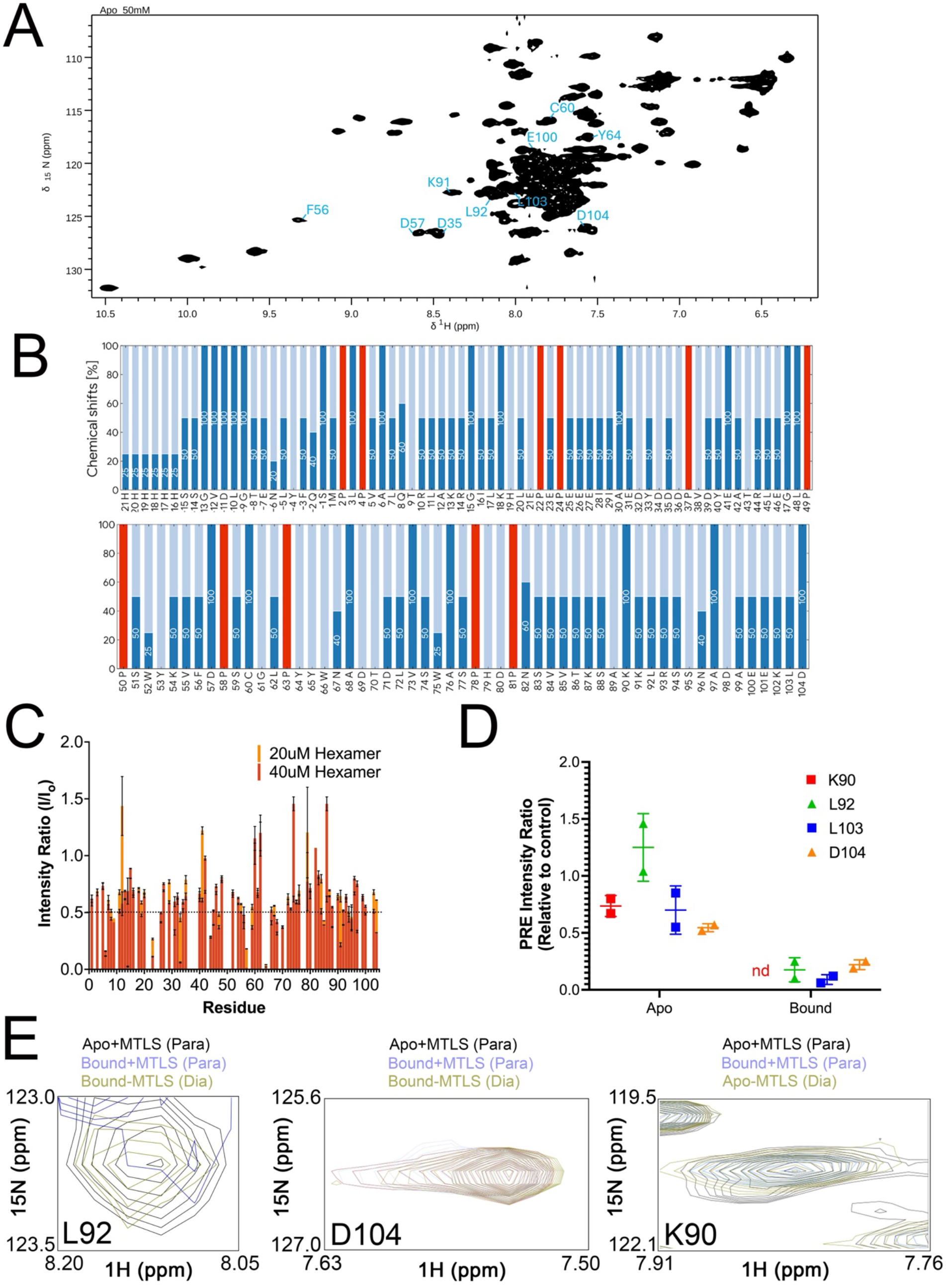
Capsid binding induces dynamics within the WW domain. (A) Backbone assignment confidence for PQBP1 (His₆-tagged construct, residues −21 to 104) generated using Bruker ARTINA, shown as percent assignment along the sequence. (B) Per-residue backbone assignment confidence for apo PQBP1 (1-104) from ARTINA. Bar height reflects confidence (More, higher %; Less, lower %). Blank positions indicate residues with no returned assignment, common for ambiguous assignments in disordered regions. Red bars mark prolines, which lack a backbone amide and cannot be assigned by amide-based NMR. (C) Per-residue HSQC intensity ratio (I/I₀) for assigned PQBP1 1-104 residues upon titration with HIV-1 capsid hexamer. PQBP1 (100 µM) was mixed with hexamer at 20 µM (1:5, yellow) or 40 µM (1:2.5, orange); values are relative to unbound PQBP1 (I₀). Dotted line indicates the 50% threshold used to define capsid-responsive residues. Error bars, SD across three independent experiments. (D) Paramagnetic relaxation enhancement (PRE) intensity ratios (I_p_/I_d_) for WW-linker residues K90, L92, L103, and D104 in the apo and hexamer-bound states (mean ± SD, n = 2 independent samples). Lower I_p_/I_d_ reflects closer proximity to the C60 spin label. nd, not determined (bound K90 signal not resolved). (E) Overlaid ¹H-¹⁵N contours for the L92, D104, and K90 amides under apo paramagnetic (Apo+MTSL, Para), bound paramagnetic (Bound+MTSL, Para), and reduced diamagnetic (Bound-MTSL or Apo-MTSL, Dia) conditions, illustrating the per-residue PRE behavior quantified in (D).

**Supplemental Figure 3:**
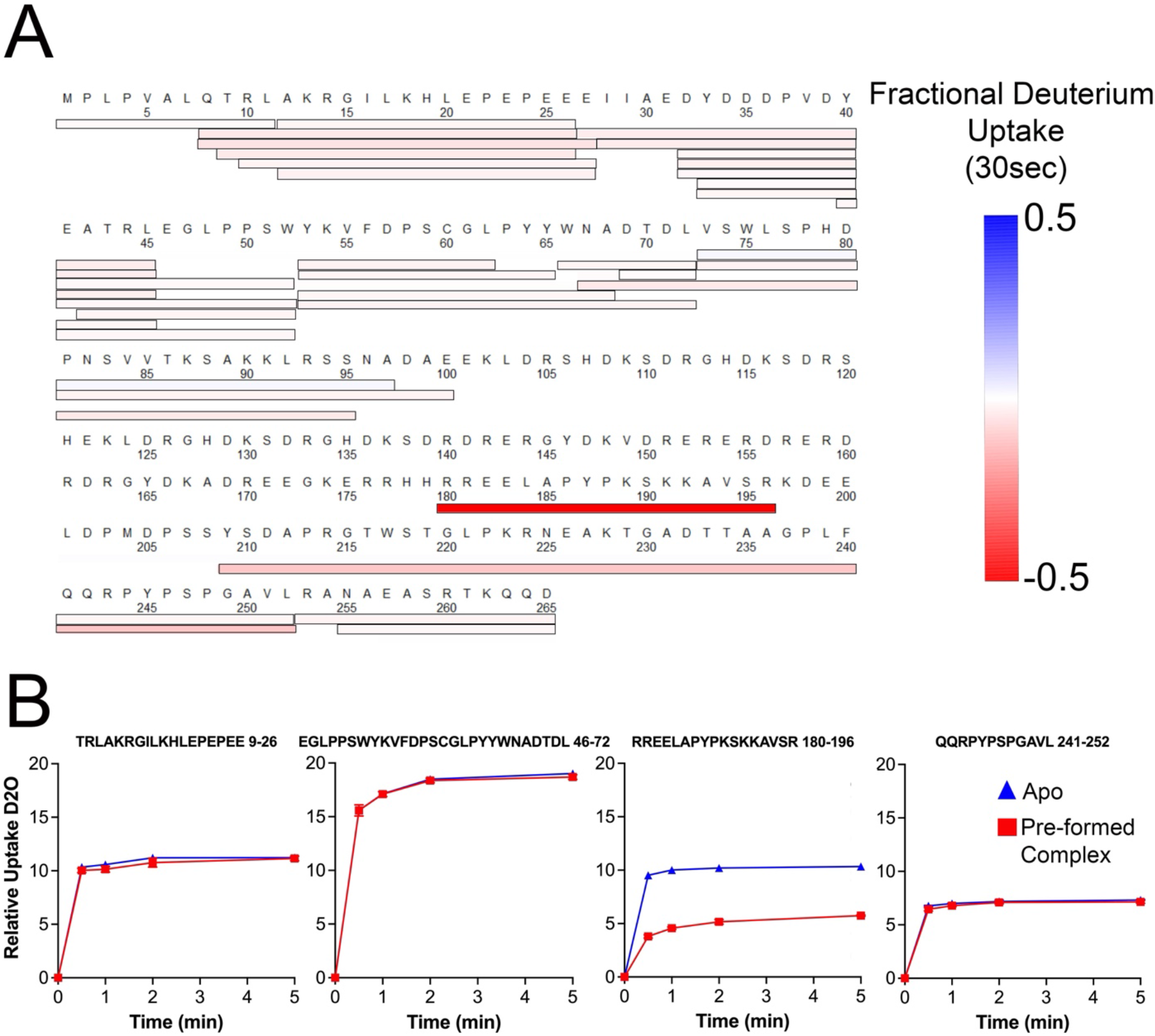
Hexamer binding promotes distal changes in PQBP1. (A) HDX-MS peptide coverage map of PQBP1 FL. Heat map of the relative fractional differential deuterium uptake differences at 30sec (ΔHDX; hexamer-bound minus unbound) mapped across the PQBP1 sequence. Regions of decreased deuterium uptake (red) indicate increased protection upon hexamer binding, while increased uptake (blue) reflects increased solvent accessibility or dynamics. (B) Relative deuterium uptake plots for PQBP1 selected peptides from each domain region (Nt, WW, PRD, and C-terminal regions) at 30 s, 1 min, 2 min, and 5 min exchange time points. Traces compare unbound and hexamer-bound conditions, with reduced uptake in the PRD indicating increased protection upon hexamer engagement. Data is representative of three independent labeling experiments (biological triplicate).

**Supplemental Figure 4:**
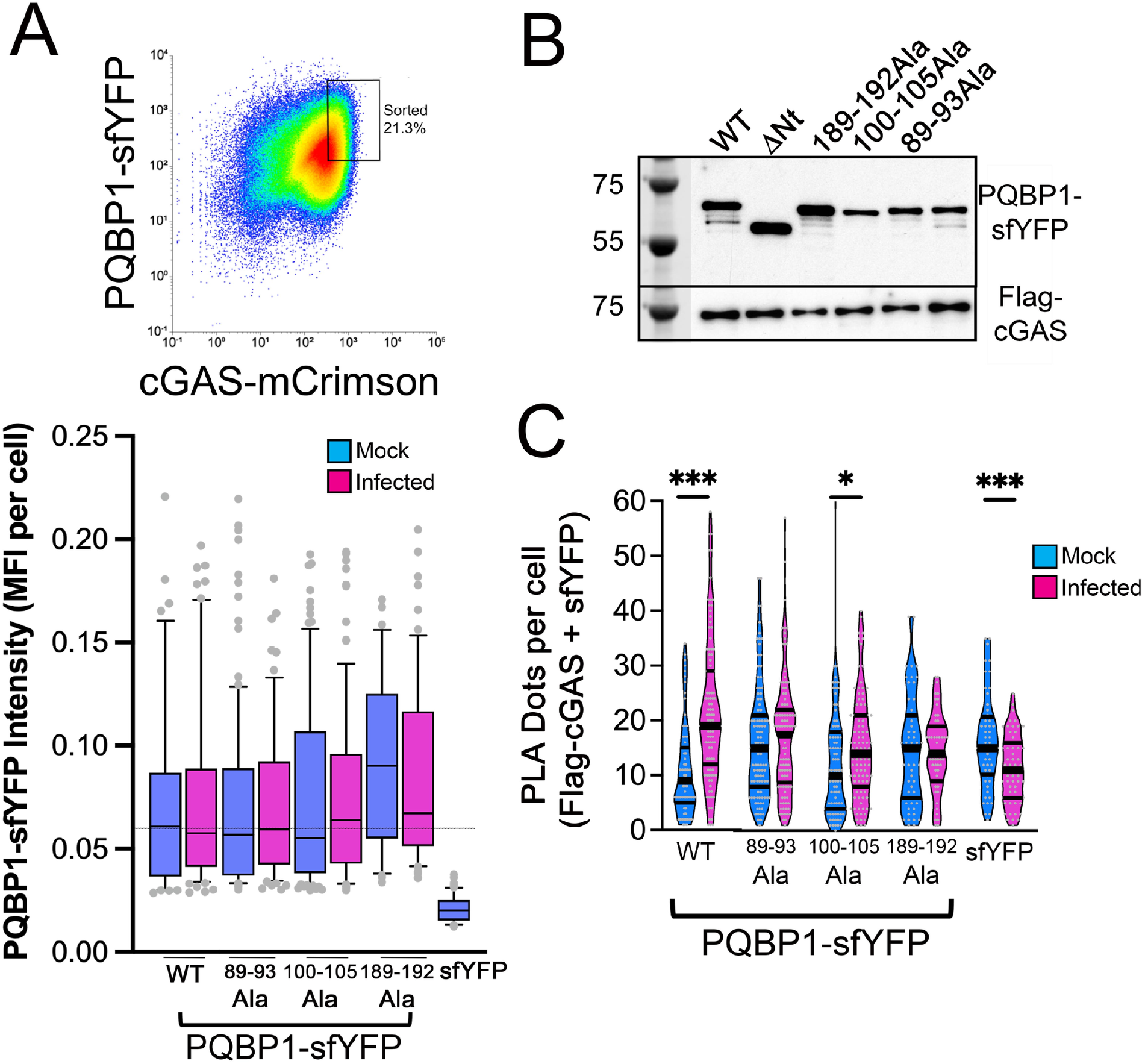
Residues within hexamer-responsive regions contribute to cGAS recruitment. (A) Representative fluorescence-activated cell sorting (FACS) plot used to isolate THP-1 cells co-expressing PQBP1-sfYFP and Flag-cGAS-mCrimson. The gated population used for downstream analyses is indicated. Mean fluorescence intensity (MFI) of PQBP1-sfYFP across cell lines used in the PLA assay. Comparable expression levels were maintained following sorting. (B) Immunoblot analysis of Flag-cGAS expression levels across cell lines used in the PLA experiments. (C) Quantification of raw PLA signal (puncta per cell) for PQBP1-sfYFP constructs under mock and HIV-1 infection conditions. Data are shown prior to normalization to illustrate absolute differences in signal across constructs from three independent experiments.

## Notes

### Competing Interest Statement

The authors have declared no competing interest.

### Summary of Updates

This version includes revisions to the main text for clarity, updated figures and supplemental files, and formatting adjustments for journal submission.

